# EpiScanpy: integrated single-cell epigenomic analysis

**DOI:** 10.1101/648097

**Authors:** Anna Danese, Maria L. Richter, David S. Fischer, Fabian J. Theis, Maria Colomé-Tatché

## Abstract

Epigenetic single-cell measurements reveal a layer of regulatory information not accessible to single-cell transcriptomics, however single-cell-omics analysis tools mainly focus on gene expression data. To address this issue, we present *epiScanpy*, a computational framework for the analysis of single-cell DNA methylation and single-cell ATAC-seq data. *EpiScanpy* makes the many existing RNA-seq workflows from *scanpy* available to large-scale single-cell data from other -omics modalities. We introduce and compare multiple feature space constructions for epigenetic data and show the feasibility of common clustering, dimension reduction and trajectory learning techniques. We benchmark *epiScanpy* by interrogating different single-cell brain mouse atlases of DNA methylation, ATAC-seq and transcriptomics. We find that differentially methylated and differentially open markers between cell clusters enrich transcriptome-based cell type labels by orthogonal epigenetic information.

## BACKGROUND

Epigenetic single-cell measurements, where the epigenetic status of single cells is evaluated using next generation sequencing techniques, are becoming mainstream^1^. Currently, two such measurements are performed routinely in the laboratory: DNA methylation status can be assessed at the single-cell level with the use of single-cell bisulfite sequencing (scBS-seq)^2^, and open chromatin patterns are investigated at individual cells using single-cell Assay for Transposase-Accessible Chromatin using sequencing (scATAC-seq)^3^. Thanks to dropping sequencing costs, well described protocols and advances in microfluidics techniques, current experimental designs afford to interrogate the epigenome of thousands of cells at the time^4–7^. These data represent a rich layer of regulatory information that stands between the genome and the transcriptome, and new analysis methods are needed to leverage it^8^.

While many methods for analyzing single-cell transcriptomics data have been developed recently^8^, this is much more limited for scATAC-seq data^9,10^ and single-cell DNA methylation data^11^, or for the joint analysis of multiple -omics data types^8^. With the current speed at which single-cell methylome and open chromatin datasets are being generated, an analysis tool that goes beyond custom-made scripts and that permits dealing with different -omics data types in the same framework is needed. Here we present *epiScanpy*, a method for the analysis of scATAC-seq and single-cell DNA methylation data, which integrates into the *scanpy* platform for single-cell transcriptomics data analysis^12^. *EpiScanpy* enables preprocessing of epigenomics data as well as downstream analyses such as clustering, manifold learning, visualization and lineage estimation. *EpiScanpy* allows for comparative analyses between -omics layers, and can serve as a framework for future single-cell multi-omics data integration. Since its downstream analyses extend the popular *scanpy* framework, it inherits properties such as fast and scalable runtime behavior and modular extensibility.

## RESULTS

### *EpiScanpy* workflow

Workflows based on *epiScanpy* consist of four stages: Feature space engineering, data pre-processing, assessment of global heterogeneity via embeddings and clusterings, and feature-level analysis to attribute drivers of heterogeneity (Fig. 1a). The input of *epiScanpy* consists of .bam files for scATAC-seq or methylation count files for single-cell DNA methylation.

**Figure 1:**
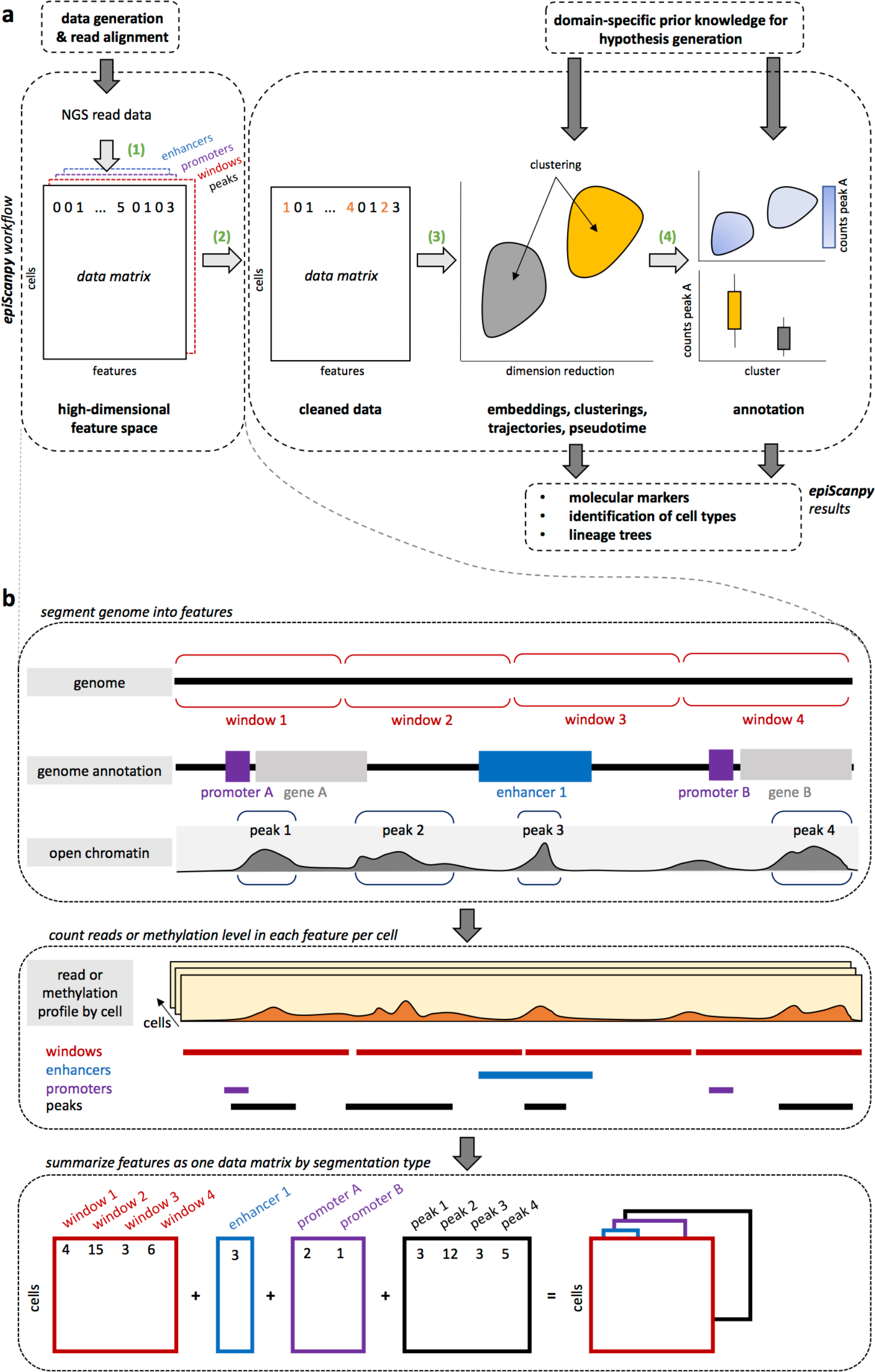
EpiScanpy workflow. **a** *epiScanpy* takes .bam files or methylation count files (for scATAC-seq and single-cell DNA methylation respectively) as input and constructs data matrices that contain read counts (for scATAC-seq) or DNA methylation levels (for single-cell DNA methylation) for different feature spaces (1). The data is pre-processed (2) and unsupervised learning algorithms (clusters, trajectories, lineage trees) are applied (3). Differential openness and methylation calling allows for cell type and lineage tree identification as well as identification of marker loci (4). **b** High dimensional feature spaces are constructed based on different genomic segmentations. The methylation level or openness per feature and per cell is calculated and summarized as a data matrix per segmentation type.

In the feature space engineering step, *epiScanpy* generates count matrices based on open chromatin levels or individual cytosine methylation levels, summarized over different sets of genomic regions (Fig. 1b). These count matrices serve as feature space that retains as much variation of the data as possible without being too high-dimensional – a feature space at single base-pair resolution can in principle be assembled but would impede downstream analysis through memory and run time issues as well as though data sparsity. These genomic regions can cover the entire genome (i.e. windows) or can be based on genomic features such as known open chromatin peaks, gene promoters, gene bodies or enhancers (suppl. methods). Any other feature space of interest, such as for example cis-regulatory topics^10^, can also be used. For scATAC-seq data, the count matrix is binarized to account for presence or absence of reads at every peak or feature, library size is regressed out and low quality single cells are filtered out (suppl. methods, Fig. SI1-2). For DNA methylation data, the CG methylation level per genomic region is computed, and features with too few covered cytosines are labelled as missing data (suppl. methods, Fig. SI3). Optionally, CH methylation can also be used for computing count matrices, but methylation in this context is only present in a limited number of mammalian tissues.

In bisulfite sequencing, it is necessary to differentiate non-methylated cytosines (zero signal) and non-observed cytosines (missing signal). Accordingly, we propose the usage of imputation methods for non-observed cytosines. Note that this is different to imputing zeros in single-cell RNA-seq, which are not inherently non-observed data points, but may also be zero count observations. *EpiScanpy* imputes based on the information from the surrounding windows or, alternatively, the population mean methylation level at the missing feature (suppl. methods). Finally, we discard non-informative features based on heuristics for the subsequent analysis: For methylation, only features which are covered in a given percentage of the cells are retained (usually ∼30%), while for ATAC-seq only the top most commonly shared peaks are considered (usually ∼20,000 peaks) (Fig. SI1-3). The constructed epigenetic data matrix is stored as an instance of the *anndata* class, a flexible data structure to store large annotated count matrices introduced in *scanpy*^*12*^. *EpiScanpy* allows for joint storage of multiple -omic modalities, allowing easy comparison between conditions and offering integrated and easy to use workflows for different types of single-cell data.

Given a processed data matrix, *epiScanpy*’s unsupervised learning algorithms can be used to uncover heterogeneity in the data, such as clusters, trajectories or lineage trees. We implemented a cell-cell distance metric based on epigenetic features to enable common algorithms that rely on a k-nearest neighbor (kNN) graph, such as Louvain clustering^13^, diffusion pseudotime^14^ and UMAP^15^. These algorithms and other unsupervised algorithms, such as tSNE^16^ and graph abstraction^17^, can directly be called via the interface to *scanpy* (Fig. 1a and suppl. methods). Note that at this point, *epiScanpy* has created an abstract representation of the data in the form of a transformed feature space or a kNN graph which can be treated in a similar fashion to single-cell RNA-seq data sets. This representation is independent of the original data form (methylation or chromatin accessibility) so that the workflows presented here truly generalize across data modalities.

Lastly, feature-level analysis dissects the drivers of heterogeneity in a data set: *epiScanpy* includes a differential methylation and differential open chromatin calling strategy (suppl. methods), which enables the ranking of genomic features (such as genes, promoters or other regulatory elements) based on their relevance in the discovered cellular identities (Fig. 1a). This allows for the identification of marker loci that can be used for a fast semi-automated cell-type identification (Fig. 1a). This feature-level analysis allows the user to correlate variation along trajectories or across clusters with marker loci to support cell type annotation and to generate hypotheses on the mechanism that underlie the identified population structure.

### Applications

To illustrate the potential of *epiScanpy* and to show how it can effectively deal with different data modalities, we applied it to brain mouse atlases from three different -omic data types: single-cell DNA methylation (snmC-seq, 3,377 prefrontal cortex neurons, 4.7% average genomic coverage^4^), single-cell open chromatin (scATAC-seq, ∼13,000 prefrontal cortex and whole brain cells, median coverage range ∼8,000 - 24,000 reads per cell^6^) and single-cell gene expression (Drop-seq, ∼690,000 cells, 9 regions of the adult mouse brain^18^).

Firstly, we explored the impact of the choice of genomic feature on the global topology (“structure”) that can be learned from the data, using clustering as an example method for unsupervised learning. Count matrices were constructed for different types of genomic features for single-cell DNA methylation and scATAC-seq data (respectively: 100kb non-overlapping windows, gene promoters, gene bodies and enhancers (from^19^); and open chromatin peaks (from^6^) and enhancers). We performed iterative Louvain clustering (suppl. methods) on each feature space and found that cells are grouped similarly across all feature spaces used, illustrating the fact that different genomic features contain partially redundant information and can be used interchangeably (Fig. 2a-c, Fig. SI4-5). For single-cell DNA methylation data, the enhancer feature space provided the clearest cell-type separation in clustering and low dimensional visualization (average silhouette score of 0.44 (enhancers) versus 0.36 (promoters), Fig.2c and Fig.2a, suppl. methods), highlighting the relevance of DNA methylation at non-genic regulatory elements at determining cell identity.

**Figure 2:**
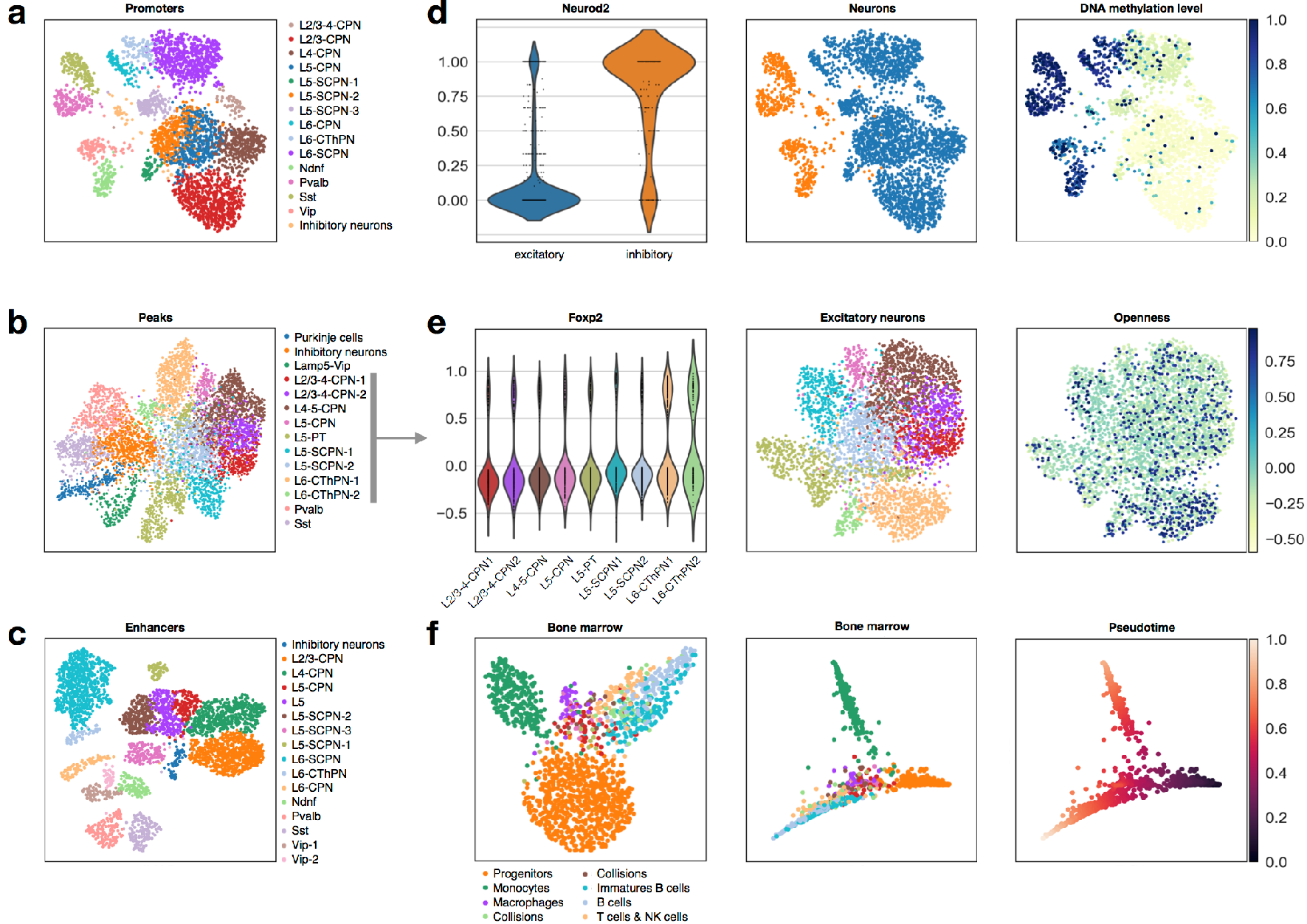
Results. **a** UMAP with Louvain clusters and annotated cell types for neurons for single-cell DNA methylation data, performed on the promoter feature space. **b** UMAP with Louvain clusters and annotated cell types for neurons for scATAC-seq data, performed on the open chromatin peak feature space. **c** UMAP with Louvain clusters and annotated cell types for neurons for single-cell DNA methylation data, performed on the enhancer feature space. **d** Differential methylation at the promoter of Neurod2 between excitatory and inhibitory neurons. **e** Differential openness at the promoter of Foxp2 in excitatory neurons. **f** UMAP (left) and pseudotime with Louvain clusters (middle) and pseudotime (right) for hematopoietic cells for scATAC-seq data.

To evaluate *epiScanpy*’s ability to map the discovered population structure in the form of clusterings to known cell types, we ranked differentially methylated and differentially open loci between the identified clusters to map cluster identity to cell types (suppl. methods). *Neurod2* was identified as one of the top differentially methylated promoters between inhibitory and excitatory neurons (Fig. 2d), which correlates with its expression levels in the adult brain^20^. *4930567H17Rik* and *Satb2* could be used to distinguish between the different neuronal layers^*20*^ and between SCPN and CPN neurons^*21*^, respectively (Fig. SI6). These observations based in CG promoter methylation are consistent with CH gene body methylation at known marker genes (Fig. SI7). Interestingly, we identify several differentially methylated promoters of genes which are not differentially expressed in the adult mouse brain^20^ but whose differential expression during embryonic development is necessary for cell fate determination, such as *Rab4a*, a marker of SST neurons expressed during E12.5 - E14.5^22^ (Fig. SI6). These findings reflect the unique ability of DNA methylation data to record past cellular states^1^ and therefore add valuable information about differentiation and lineage trees to models based on transcriptomics. This integration of complementary layers of information highlight the potential of multi-omics approaches to build a more complete picture of developmental systems.

For scATAC-seq, we identified top differentially open peaks which were used to label cell clusters (Fig. 2b). For example, openness of the *Ndrg2* promoter can be used to distinguish astrocytes^23^ (Fig. SI8) and microglia and oligodendrocytes are identified by open peaks in the promoters of *Runx1*, and *Efnb3* respectively^24,25^ (Fig. SI8). As a whole group, neurons show openness of peaks in promoters for neuronal genes like *Ptprd, Pik3r1*, and *Syt1*^*26–28*^ (Fig. SI8), while differential openness at the *Foxp2* promoter can be used to identify Layer 6 cortical neurons (Fig. 2e), for example. A comprehensive list of differential markers used for single-cell DNA methylation and scATAC-seq cluster identification can be found in SI Table 1.

We compared *epiScanpy* cell type identification to the one provided by Luo *et al.* (obtained using CH-gene-body methylation levels)^4^ and the one provided by Cusanovich *et al.* (obtained using promoter and distal regulatory site accessibility)^6^. Respectively ∼89% and ∼71% of cells are assigned to the same cell type as in the original publications (Fig. SI9-10). For the scATAC-seq dataset the biggest discrepancy is found between SCPN/CPN assignments, where we identify clusters with SCPN signatures that were labelled as CPN neurons in the original publication, and vice versa (Fig. SI11).

We performed a global comparison of multi-omic cellular atlases based on mouse brain tissue from single-cell DNA methylation, scATAC-seq and scRNA-seq data (processed using *scanpy*, suppl. methods). While some markers are differentially expressed, differentially open and differentially methylated between clusters (Fig. SI12), there is also a large number of non-redundant markers, such as that of *Fabp7. Fabp7* is a brain fatty acid binding protein that has been reported to be important for forebrain physiology and is associated with Schizophrenia^29^, which displays signs of differential regulation in CPN neurons (differentially open and methylated) but is not expressed in neurons (Fig. SI12). These markers provide complementary information between data modalities, underpinning the fact that every -omic layer contributes its individual non-redundant layer of information, and emphasizing the need for a tool that deals with many -omic data types and facilitates integration across modalities.

Finally, we also considered open chromatin profiles of hematopoietic cells (bone marrow cell types from^6^) to evaluate whether *epiScanpy* can learn developmental trajectories with pseudotime and more complex lineage trees with graph abstraction directly based on epigenomic profiles (Fig. 2f). Such continuous descriptions of developmental systems have been very useful in studies based on single-cell transcriptomics. *EpiScanpy* discovers 7 cell types (Fig. SI13) and recovers the known hematopoietic differentiation tree (Fig. 2f).

## DISCUSSION

In summary, *epiScanpy* is a fast and versatile tool for the analysis of single-cell epigenomic data and its integration with single-cell transcriptomic data. It offers the first unified framework for the analysis of both single-cell DNA methylation, scATAC-seq and single-cell transcriptomic data, and its flexible data structure is ready to handle other new types of single-cell -omic data, such as Hi-C or NOME-seq, as well as multi-omics single-cell data. *EpiScanpy* addresses the open question of feature space construction on epigenetic data and we show evidence that similar manifolds can be learned based on different feature spaces. *EpiScanpy* also scales well to the large scATAC-seq data sets generated with the 10x Chromium platform (Fig. SI14)^30^. *EpiScanpy* performs single-cell graph construction from potentially any type of single-cell -omics data and performs downstream analysis like low-dimensional data visualization, clustering, single-cell graph abstraction or trajectory inference, and differential calling. *EpiScanpy* is available as a python package through Github (https://github.com/colomemaria/epiScanpy, documentation available on episcanpy.readthedocs.io) and builds upon the *scanpy* analysis toolbox^12^, opening the toolchain to the commonly measured single-cell epigenomic data.

## ACKNOWLEDGMENTS

We would like to thank Boyan Bonev for discussions on brain cell type identification, as well as Alex Wolf and Philip Angerer for consultation on integration into *scanpy* and for sharing demultiplexing scripts. We would like to thank Meshal Ansari for her input on the usage of dimension reduction techniques for scATAC-seq.

This work was supported by the Impuls-und Vernetzungsfonds of the Helmholtz-Gemeinschaft (grant VH-NG-1219) for M.C.T. F.J.T. acknowledges financial support by the German Science Foundation (SFB 1243 and Graduate School QBM) as well as the Federal Ministry of Education and Research (Single Cell Genomics Network Germany). A.D. was supported by the Incubator grant sparse2big (grant #ZT-I-0007). D.S.F. acknowledges financial support by a German research foundation (DFG) fellowship through the Graduate School of Quantitative Biosciences Munich (QBM) (GSC 1006) and by the Joachim Herz Stiftung.

## Contributions

M.C.T. and F.J.T. designed the study. A.D., M.L.R. and D.F. developed the method. A.D. and M.L.R. analysed data. M.C.T., A.D. and D.F. wrote the manuscript.

## Code availability

*EpiScanpy* is available through Github (https://github.com/colomemaria/epiScanpy) and the documentation is available on episcanpy.readthedocs.io.

## Data availability

The data sets analysed here were downloaded from GEO (see supplementary text for GEO accession codes).

