## Supplemental text for "EpiScanpy: integrated single-cell epigenomic analysis"

**Contents :**

**DATASETS** 3

**ANNOTATIONS** 3

**CHROMATIN ACCESSIBILITY** 4

scATAC-seq Peak processing 4

Enhancers based count matrix 4

Cell type identification and comparison with Cusanovich et al. 4

Bone marrow processing and diffusion pseudotime 5

**METHYLATION PROFILES** 5

snmC-seq data processing 5

IMPUTATION OF MISSING VALUES - Methylation 6

Cell type identification 7

**scRNA-seq PROCESSING** 7

**ATLAS COMPARISON** 8

**DATASETS**

The single cell DNA methylation, scATAC-seq and scRNA-seq data sets have been described in Luo et al.[1], Cusanovich et al.[2] and Saunders et al.[3], respectively.

The mouse **frontal cortex single cell transcriptome** from Saunders et al.[3] data are available on GEO: GSE116470. Saunders et al. measured gene expression in ~690,000 cells from 9 regions of P60 C57BL/6 male mice brain using Drop-seq , and reads were aligned on GRCm38.81 assembly of the mouse genome.

The **methylation profiles** from Luo et al.[1] were downloaded from GEO: GSE97179. This sc DNA methylation dataset was obtained applying single-nucleus methylcytosine sequencing (snmC-seq) to 3377 prefrontal cortex neurons (NeuN+) of 8 week old mice, yielding 4.7% average coverage of the mouse genome per single cell. The different neuron subtypes were not labelled experimentally. The downloaded datasets contain cytosine summaries for all cells, with columns for genomic position, number of methylated cytosines, number of total cytosines, and cytosine context (CG, CH; where H=A,T,C). The sequencing data were aligned on GRCm38, all processing specifications and metadata are available as supplementary material in Luo et al.[1].

For the **chromatin accessibility** dataset, we downloaded whole brain, prefrontal cortex and bone marrow data (replicate 62216, ~13,000 brain cells, <7000 bone marrow cells, median coverage range of ~8,000 - 24,000 reads per cell, aligned on MGSCv37) generated by Cusanovich et al.[2] as raw binary data matrices (peaks and windows) from NCBI, GEO: GSE111586. Cell specific metadata were downloaded from http://atlas.gs.washington.edu/mouse-atac/data/. Raw bam files per replicate for scATAC-seq were also downloaded from http://atlas.gs.washington.edu/mouse-atac/data/.

**ANNOTATIONS**

*Promoter annotations*: TSS positions from the Eukaryotic Promoter Database[4] were used, and promoters were defined as -1.5kb upstream of the TSS and +500 bp downstream.

*Enhancer annotations*: enhancer annotation was downloaded from the Mouse Encode Project at Ren Lab[5]. Enhancers are 2000 bp long. The mm9 annotations were converted to mm10 using LiftOver[6].

*Gene body annotations*: gene bodies from UCSC (mm10) were used.

*Peaks*: to find peaks in promoters, the list provided by Cusanovich et al.[2], was taken http://krishna.gs.washington.edu/content/members/ajh24/mouse atlas data release/metadata/peak promoter intersections.txt

### CHROMATIN ACCESSIBILITY

#### scATAC-seq Peak processing

The whole brain and prefrontal cortex replicates were merged, yielding an initial raw matrix of 11453 cells and 436206 peaks (matrix_1). Matrix_1 was reduced to only contain peaks shared in at least 1000 cells, generating a new matrix with the top 19535 most covered peaks (matrix_2). Afterwards, we regressed the total number of peaks covered per cell from matrix_1 on matrix_2. After linear regression, cells containing less than 600 open peaks were discarded. The final processed matrix has 11436 cells and 19535 peaks and was saved as an *anndata* object.

Dimensionality was reduced by calculating principal components (PCs). We constructed a nearest neighbor graph using Euclidean distance between cells based on feature openness. Cell clusters were defined using Louvain clustering method[7].

We identified a very distinct dataset, exclusively coming from the whole brain replicate, corresponding to Cerebellar granular cells, that we excluded from later analysis.

#### Enhancers based count matrix

In order to quantify enhancer openness, the prefrontal cortex sample .bam file was demultiplexed using a custom perl script. Then, the enhancer count matrix was created from demultiplexed .bam files using epiScanpy and saved as an *anndata* object. Afterwards the matrix was binarized to account for the presence/absence of reads, and enhancers covered in less than 100 cells were removed, resulting in a final set of 10,505 enhancers. Additionally, cells that contained less than 200 covered enhancers were removed, reducing the cell number from initially 5,958 cells to 3,670 cells. Finally, the total number of peaks from the unfiltered enhancer matrix was regressed out on the filtered one.

#### Cell type identification and comparison with Cusanovich et al.

Cell types were defined based on openness in promoters for known marker genes ( supplementary table 1) and these annotations were compared to the ones provided by Cusanovich et al.[2]. We first identified broad cell types (Astrocytes, Oligodendrocytes, Microglia, Endothelial cells, Excitatory neurons, Inhibitory neurons) as well as collision cells, using known marker genes (supplementary table 1). Then we refined our cell clustering to identify neuron subtypes. This was done by considering only neurons and splitting them initially into two clusters, corresponding to excitatory and inhibitory neurons. These two clusters were split into further subclusters. Using multiple pairwise t-tests with multiple testing correction (Benjamini-Hochberg) we identified, for each neuron subcluster, the top 400 most differential peaks (p-value <0.05) versus the rest of the neurons, and versus the rest of the inhibitory only and excitatory only neurons.  We obtained the genes corresponding to these peaks using the overlapping promoter annotations (data from Cusanovich et al.[2] http://krishna.gs.washington.edu/content/members/ajh24/mouse). To identify neuronal subtypes according to the layers (2/3, 4, 5, 6) and the projections (PT, CPN, SCPN and CThPN), we extracted a list of top marker genes from the Allen brain atlas[8]. Finally, the list of most differentially open genes was crossed with these marker genes to find the most likely neuron type for every cluster (supplementary table 1).

We compared the cell annotation obtained using epiScanpy to the cell annotation from Cusanovich et al.[2]. The number of cells falling into each annotation was counted and normalized by the total number of cells per group and the normalized counts were plotted as heatmaps (SI fig 9).

For the analysis based on enhancer openness, the original cell type annotation from Cusanovich et al.[2], as well as the cell annotation obtained with epiScanpy from the peak count matrix, were used to identify cell types.

**Bone marrow processing and diffusion pseudotime**

The raw binary peak matrix was loaded as *anndata* object, peaks covered in less than 500 cells (out of 6949 cells) and cells with less than 500 peaks covered were removed. We obtained a count matrix with 2710 cells and 20001 peaks, the total number of peaks per cell were regressed out.

Then, we built a neighborhood graph using Euclidean distance, low dimensional representation and Louvain clustering to identify the major expected cell types. We compared our clusters to Cusanovich et al.[2] annotation to validate and label our clusters.

Finally, we excluded cells labeled as erythroblast to focus on less abundant and differentiating cells.

To look at the dynamical changes of the chromatin landscape upon differentiation, we computed diffusion pseudotime and diffusion map for a better low dimensional representation (using the default scanpy implementation). We defined the hematopoietic progenitor cluster as the root cell and inferred cell progression along the graph using geodesic distance (Fig. 2e and Fig. SI13).

### METHYLATION PROFILES

#### snmC-seq data processing

Starting from the methylation summary files obtained from Luo et al.[1], we built count matrices based on different segmentations of the genome. We used epiScanpy to build count matrices with 100kb non overlapping windows, promoters, gene bodies and enhancers.

For each feature and each cell of the count matrix, we computed the average methylation level based on the number of methylated and unmethylated reads of cytosines either in a CG or CH (H=A, T, C) context. For large genomic regions such as 100kb windows, we set a minimum number of cytosines (in CG or CH context) to be covered in order to calculate an average methylation level. If these minima are not reached, a 'NA' (Not Available) is added in the matrix. For smaller genomic regions such as promoters or enhancers the minimum number of covered cytosines (in CG or CH context) is set to one, otherwise the features is labelled as missing data ('NA') (we tested higher cutoffs and the performance did not change).

The raw count matrices contain a large number of missing values. We filter out features with insufficient number of covered cells and impute the missing information for the remaining missing features/cells (cf. imputation of missing values). For each of our count matrices, we filtered out features shared in less than a 1000 cells and removed cells with less than a 1000 features covered before imputation.

**IMPUTATION OF MISSING VALUES - Methylation**

Since the single cell DNA methylation dataset from Luo et al. has a genomic coverage of ~4.7% on average[1], methylation count matrices are very sparse, containing many missing values that correspond to uncovered cytosines and which are represented as 'NA'. Depending of the size of the features considered to build the count matrix, as well as the minimum coverage threshold required, the distribution of the missing values change drastically (Fig. SI3). In order to perform dimensionality reduction, we filter cytosines which are not covered in the majority of cells, and for the remaining ones we impute missing information in the uncovered cells.

Two different imputation approaches are available on epiScanpy. The first approach, which is the default one, is to calculate the average methylation level at every feature after filtering, and assign this average value at the missing cells. After clustering it is possible to refine the imputed value by assigning the average methylation level of the cells within a cluster. A second imputation approach is to load an additional count matrix for larger features than the matrix of interest (for example, 50kb window count matrix versus promoter count matrix) and use the information from the larger feature to impute the missing value of the overlapping smaller feature. In the case where the overlapping feature is also missing, it is possible to extend the it to the adjacent feature. After that, features that are still 'NA' are either discarded or imputed based on the first approach. An alternative filtering option available on epiScanpy is to filter the features not based on the number of cells covered but on the number of different methylation level it contains. This approach will retain lowly covered but highly variable features while discarding well covered but very uniformly methylated features.

**Cell type identification**

For all imputed count matrices corresponding to the different annotations, we computed principal components and ratio of explained variances. Then we built a neighborhood graph using Euclidean distance. For low dimensional representation we used principal component analysis (PCA), t-distributed stochastic neighborhood embedding (t-SNE) and uniform manifold approximation and projection (UMAP)[9].

We checked for potential experimental biases and batch effect (plates, library preparations) and did not find any significant effects. Additionally, we looked at two important biological sources of variation that might affect the quality of the clustering. First, we regressed out the overall methylation level of the cell. Second, we set the average methylation level of each feature to one by diving every feature methylation levels by its average feature methylation across all cells. The CG methylation level ranges from 0 to 1 within the different data matrices, representing an important variation between the different variables. In both cases, normalising theses variables did not improve our clustering.

We used Louvain clustering on every feature matrix independently (i.e. windows, promoters, enhancers and gene bodies), to identify initially two main clusters, corresponding to excitatory and inhibitory neurons. We used multiple pairwise Welch's t-test with multiple testing correction (Benjamini-Hochberg) to obtain the most differentially methylated features between these clusters. Using the promoter count matrix for cell type identification, we obtained the top 400 most differentially methylated promoters between the two clusters. Among the top results we identified promoters of known marker genes for both excitatory and inhibitory neurons and labelled the clusters accordingly. We repeated the clustering and cell type identification iteratively treating inhibitory and excitatory neurons independently, by dividing the matrix into two submatrices.

As a first validation of the obtained promoter CG methylation based cell type identification, we used CH gene body methylation of known marker genes (Fig. SI7). We also compared our cell type annotation to the one obtained by Luo et al.[1] obtained by the authors using solely CH methylation, as a second validation (Fig. SI9).

Finally, we computed silhouette scores for promoter- and enhancer-based analysis using the scikit-learn function silhouette[10], which has been implemented as part of *epiScanpy*.

**scRNA-seq PROCESSING**

The raw count matrix as well as the metadata (https://storage.googleapis.com/dropviz-downloads/static/regions/F_GRCm38.81.P60Cortex_noRep5_FRONTALonly.raw.dge.txt.gz) were converted into *anndata* format for further processing using scanpy[11]]. Genes not covered in a minimum of 30 cells and cells that had less than 200 genes covered or with more than 15 percent mitochondrial reads were filtered out. Data were normalized using linear regression using the total number of reads per cell.

**ATLAS COMPARISON**

As the methylation data only consists of neurons, we limited scRNA-seq and scATACseq matrices to identified neuron clusters for comparison. Additionally, the number of cells and features differ between the three datasets, and not all features overlap (e.g. not all genes are present in the expression matrix, not all promoters are considered in the methylation count matrix, and not all peaks fall into genes). Hence, for each -omic layer (methylation, open chromatin and transcriptome separately), we identified the top 400 most differentially methylated promoters, most differentially open peaks (then restricted to peaks in promoters) and most differentially expressed genes between clusters (with p-val < 0.05). We obtained 3365 methylated promoters, 1714 peaks (in promoters) and 2285 transcripts. If we extended the analysis to the top 800 most differential features, we obtained 5601 promoters, 3657 peaks, and 2601 genes.

| | **Top 400 markers** | **Top 800 markers** |
|---|---|---|
| Unique to methylation | 2735 | 4038 |
| Unique to ATAC | 1267 | 2395 |
| Unique to RNA | 1737 | 1597 |
| ATAC + methylation | 288 | 982 |
| RNA + methylation | 389 | 724 |
| RNA + ATAC | 206 | 423 |
| RNA + ATAC + methylation | 47 | 143 |

Of these, most features are unique to one omic and only a few genes/promoters are differential across all omic layers. The largest number of shared features between two omic layers is between open chromatin and methylation. Two main reasons likely explain these limited overlaps. One is that every omic layer contain specific information. The second reason is the very different number of features that can possibly be measured in the different omic layers. Even restricted to coding regions of the genome, DNA based measurements (open chromatin and methylation) can provide information that is not present in expression, thus providing different insights on the cell identity and potential.


1. Luo, C. *et al.* Single-cell methylomes identify neuronal subtypes and regulatory elements in mammalian cortex. *Science* **357**, 600–604 (2017).
2. Cusanovich, D. A. *et al.* A Single-Cell Atlas of In Vivo Mammalian Chromatin Accessibility. *Cell* **174**, 1309–1324.e18 (2018).
3. Saunders, A. *et al.* Molecular Diversity and Specializations among the Cells of the Adult Mouse Brain. *Cell* **174**, 1015–1030.e16 (2018).
4. Dreos, R., Ambrosini, G., Groux, R., Cavin Périer, R. & Bucher, P. The eukaryotic promoter database in its 30th year: focus on non-vertebrate organisms. *Nucleic Acids Res.* **45**, D51–D55 (2017).
5. Shen, Y. *et al.* A map of the cis-regulatory sequences in the mouse genome. *Nature* **488**, 116–120 (2012).
6. Hinrichs, A. S. *et al.* The UCSC Genome Browser Database: update 2006. *Nucleic Acids Res.* **34**, D590–8 (2006).
7. Blondel, V. D., Guillaume, J.-L., Lambiotte, R. & Lefebvre, E. Fast unfolding of communities in large networks. *J. Stat. Mech.* **2008**, P10008 (2008).
8. Staining, H. Allen cell types database. *Allen Inst. Brain Sci* **88**, 1–9 (2015).
9. McInnes, L., Healy, J. & Melville, J. UMAP: Uniform Manifold Approximation and Projection for Dimension Reduction. (2018).
10. Pedregosa, F. *et al.* Scikit-learn: Machine Learning in Python. *J. Mach. Learn. Res.* **12**, 2825–2830 (2011).
11. Wolf, F. A., Angerer, P. & Theis, F. J. SCANPY: large-scale single-cell gene expression data analysis. *Genome Biol.* **19**, 15 (2018).
