## Supplemental figures for "EpiScanpy: integrated single-cell epigenomic analysis"

**Affiliations:**
[1] Institute of Computational Biology, Helmholtz Center Munich, German Research Center for Environmental Health, Neuherberg, Germany
[2] European Research Institute for the Biology of Ageing, University of Groningen, University Medical Centre Groningen, Groningen, The Netherlands
[3] TUM School of Life Sciences Weihenstephan, Technical University of Munich, Freising, Germany
[4] Department of Mathematics, Technical University of Munich, Garching, Germany

* correspondence to:


**Suppl. Figure 1: scATAC-seq peak commonness**

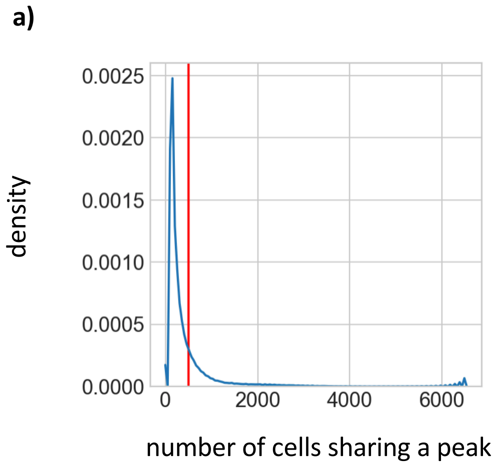


**Suppl. Figure 1: a)** Density distribution of peak commonness (scATAC-seq) between all cells (cells in “prefrontal cortex” and “whole brain”). The line in red indicates the cutoff chosen to reduce feature space to only consider peaks that are shared in at least 500 cells. **b)** Histogram of the number of peaks per cell.

**Suppl. Figure 2: scATAC-seq processing steps**

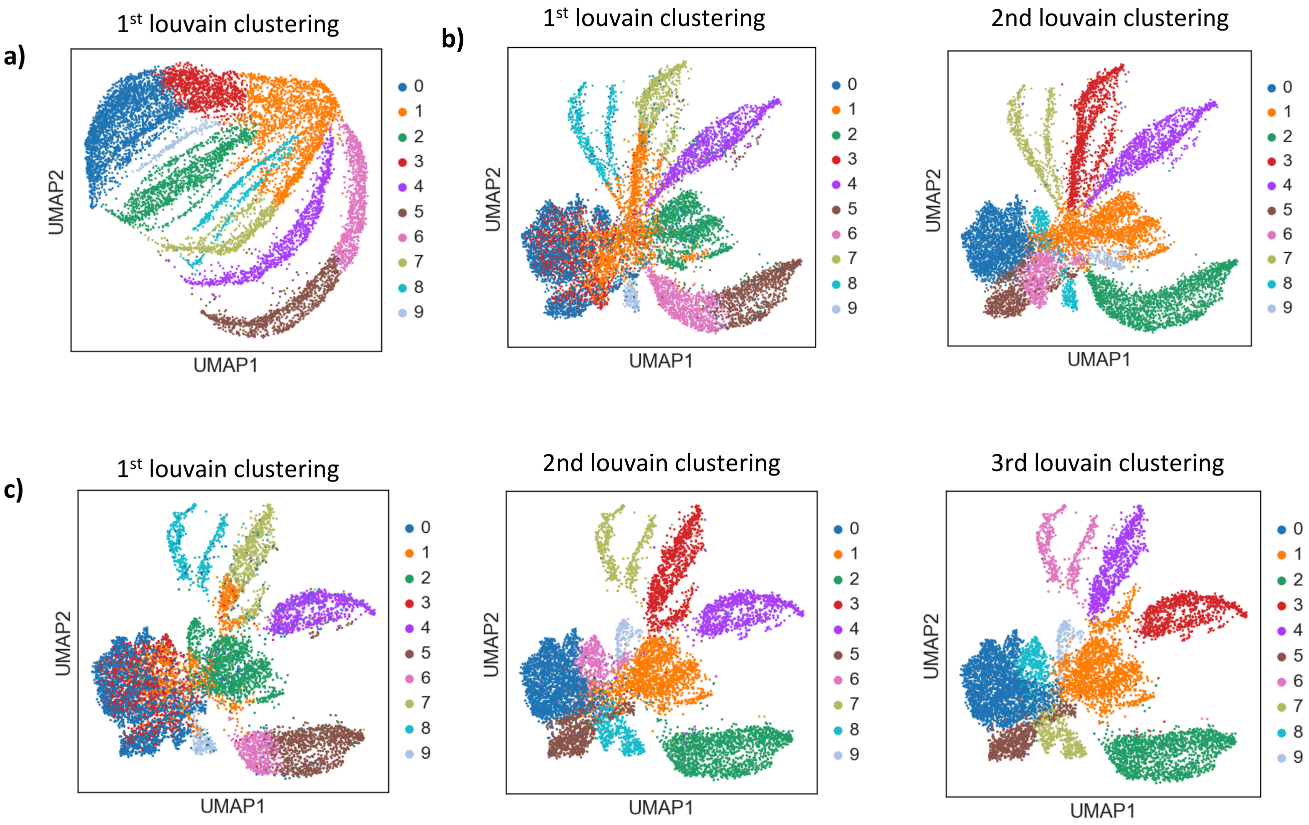


**Suppl. Figure 2: a)** UMAP visualization of the raw scATAC-seq data with initial Louvain clustering (11453 cells × 19535 peaks) **b)** UMAP visualization of the scATAC-seq data after regressing out the number of peaks per cell, with the same Louvain cluster labels as in a) **(left)** and with second Louvain cluster labels obtained with the new regressed dataset **(right)**. **c)** UMAP visualization of the scATAC-seq data after regressing out the number of peaks per cell and removing lowly covered cells (cells with less than 600 peaks covered), with **(from left to right)** the 1st Louvain clusters as in a), the 2nd set of Louvain clusters and finally new Louvain clusters obtained with the regressed dataset after removing lowly covered cells (9513 cells × 19535 peaks). These plots highlight the importance of data preprocessing in scATAC-seq.

**Suppl. Figure 3: sc DNA methylation feature commonness**

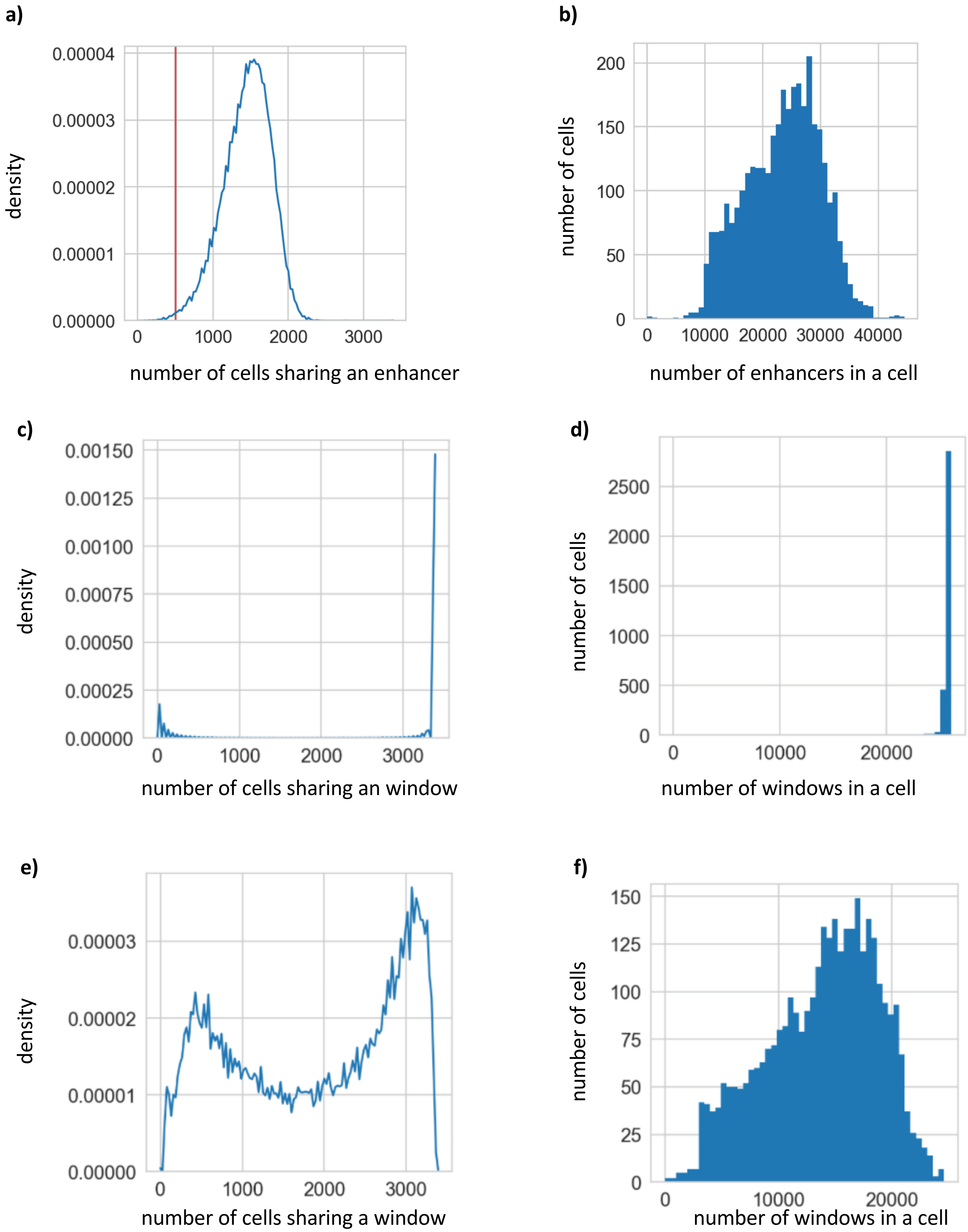

**Suppl. Figure 3 (continued)**

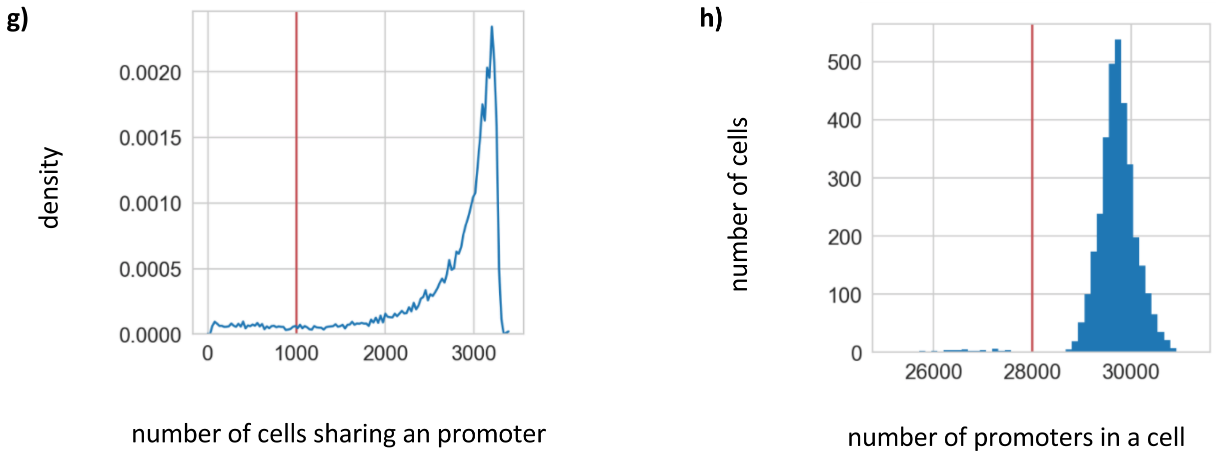


**Suppl. Figure 3: a)** Density distribution of enhancer commonness (DNA methylation) between all cells, for enhancers with at least 1 covered cytosine in CG context. The red line indicates the cutoff chosen to reduce feature space to only consider enhancers that are shared in at least 500 cells. **b)** Histogram of the number of enhancers per cell, with at least 1 CG covered. **c)** Density distribution of 100 kb window commonness (DNA methylation) between all cells, for windows with at least 1 covered cytosine in CG context. The red line indicates the cutoff chosen to reduce feature space to only consider windows that are shared in at least 500 cells. **d)** Histogram of the number of windows per cell, with at least 1 CG covered. **e), f)** Same as c) and d) but with windows with at least 30 covered CGs. **g)** Density distribution of promoter commonness (DNA methylation) between all cells, for promoters with at least 1 covered cytosine in CG context. The red line indicates the cutoff chosen to reduce feature space to only consider promoters that are shared in at least 1000 cells. **h)** Histogram of the number of promoters per cell, with at least 1 CG covered. The line in red indicates the cutoff chosen to discard cells which do not have enough covered promoters (above 28000 promoters).

**Suppl. Figure 4: scDna methylation clustering with different features**

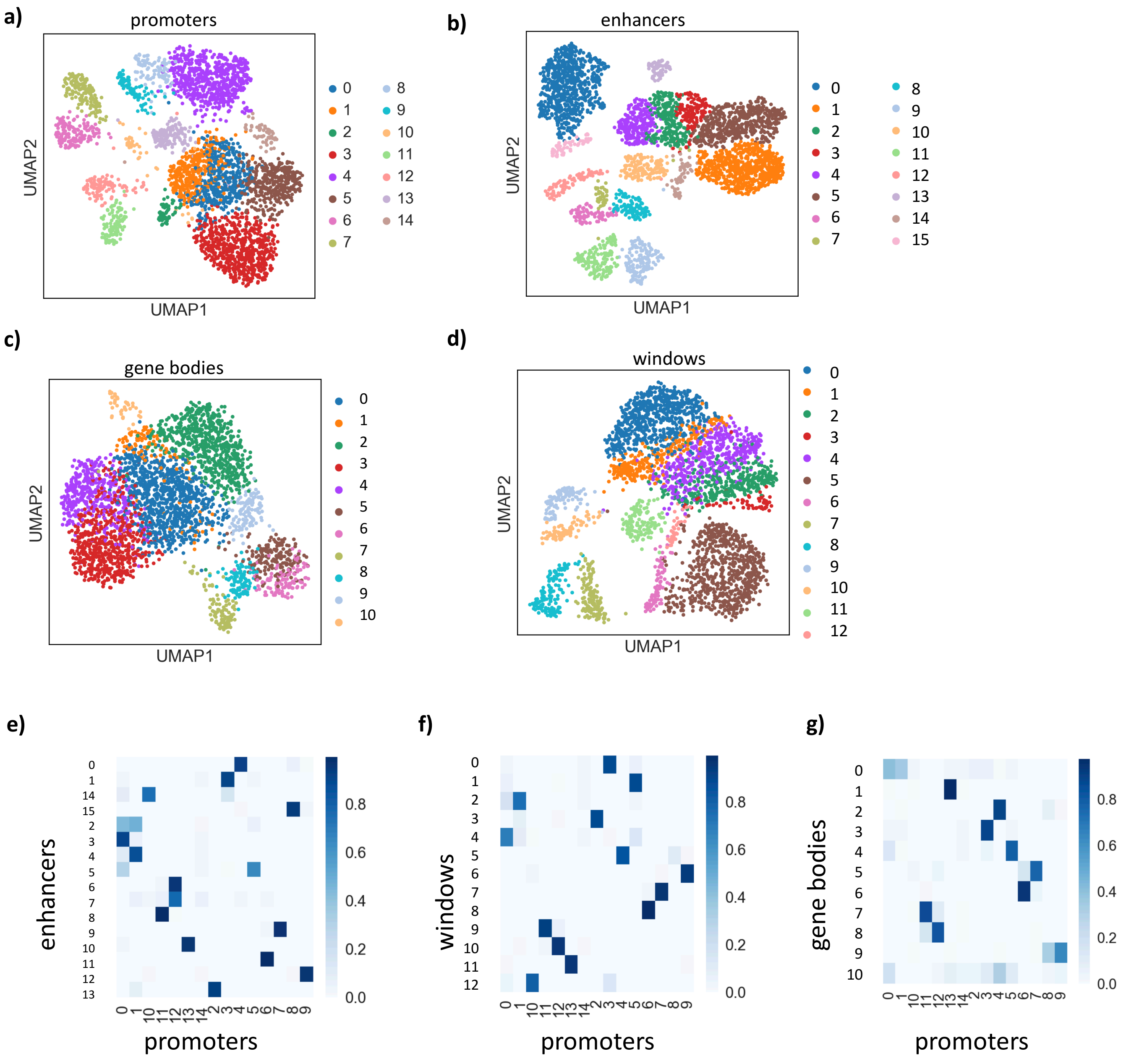


**Suppl. Figure 4: a)** sc DNA methylation clustering for prefrontal cortex neurons using different feature spaces: **a)** promoters CG methylation (3321 cells x 27393 promoters), **b)** enhancer CG methylation ( 3358 x 50018 enhancers), **c)** gene body CG methylation (3233 cells x 26947 gene bodies), **d)** 100kb windows (3224 cells x 25236 windows). **e-g)** heatmap for cluster identification comparison between **e)** promoters and enhancers, **f)** promoter and windows and **g)** promoters and gene bodies.

**Suppl. Figure 5: scATAC-seq clustering with different features**

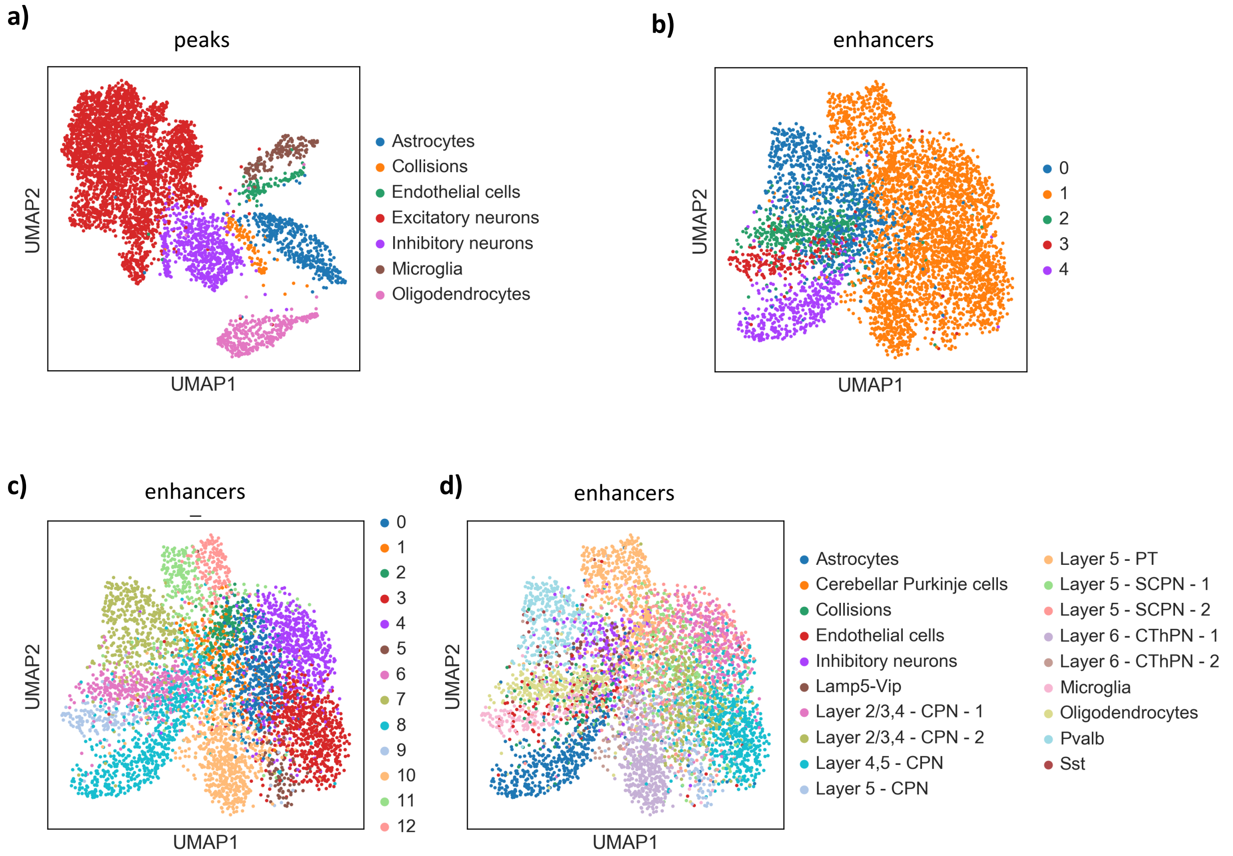


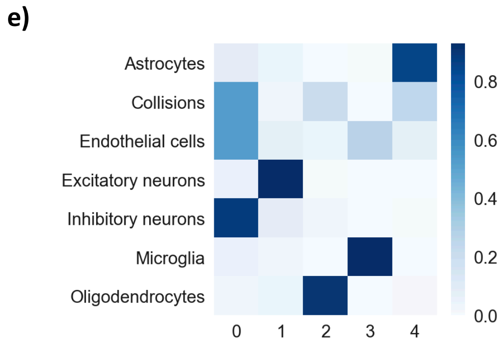


f)

| | 0 | 1 | 2 | 3 | 4 |
|---|---|---|---|---|---|
| **Astrocytes** | 37 | 17 | 3 | 4 | 357 |
| **Collisions** | 47 | 2 | 19 | 0 | 22 |
| **Endothelial cells** | 51 | 8 | 4 | 26 | 7 |
| **Excitatory neurons** | 191 | 3054 | 31 | 10 | 5 |
| **Inhibitory neurons** | 554 | 56 | 13 | 2 | 5 |
| **Microglia** | 8 | 4 | 0 | 157 | 0 |
| **Oligodendrocytes** | 13 | 15 | 313 | 2 | 4 |

**Suppl. Figure 5: : a)** scATAC-seq clustering for prefrontal cortex cells for the peak features (5041 cells × 19535 peaks), with the annotation as described in the main text. **b)** scATAC-seq clustering for prefrontal cortex cells for enhancer features (5958cells × 10505 enhancers), with five Louvain clusters. **c)** scATAC-seq clustering for prefrontal cortex cells for enhancer features (5958cells × 10505 enhancers), with 13 Louvain clusters. **d)** scATAC-seq clustering for prefrontal cortex cells for enhancer features (5958cells × 10505 enhancers), with cluster color as described in the main text for peak features (main text figure 2c). **e)** Heatmap comparing cell type identification between peak and enhancer features (a) and b)), with peaks per row and enhancers per column. **f)** Number of cells per cluster (with peaks per row and enhancers per column).

**Suppl. Figure 6: scDNA methyation**

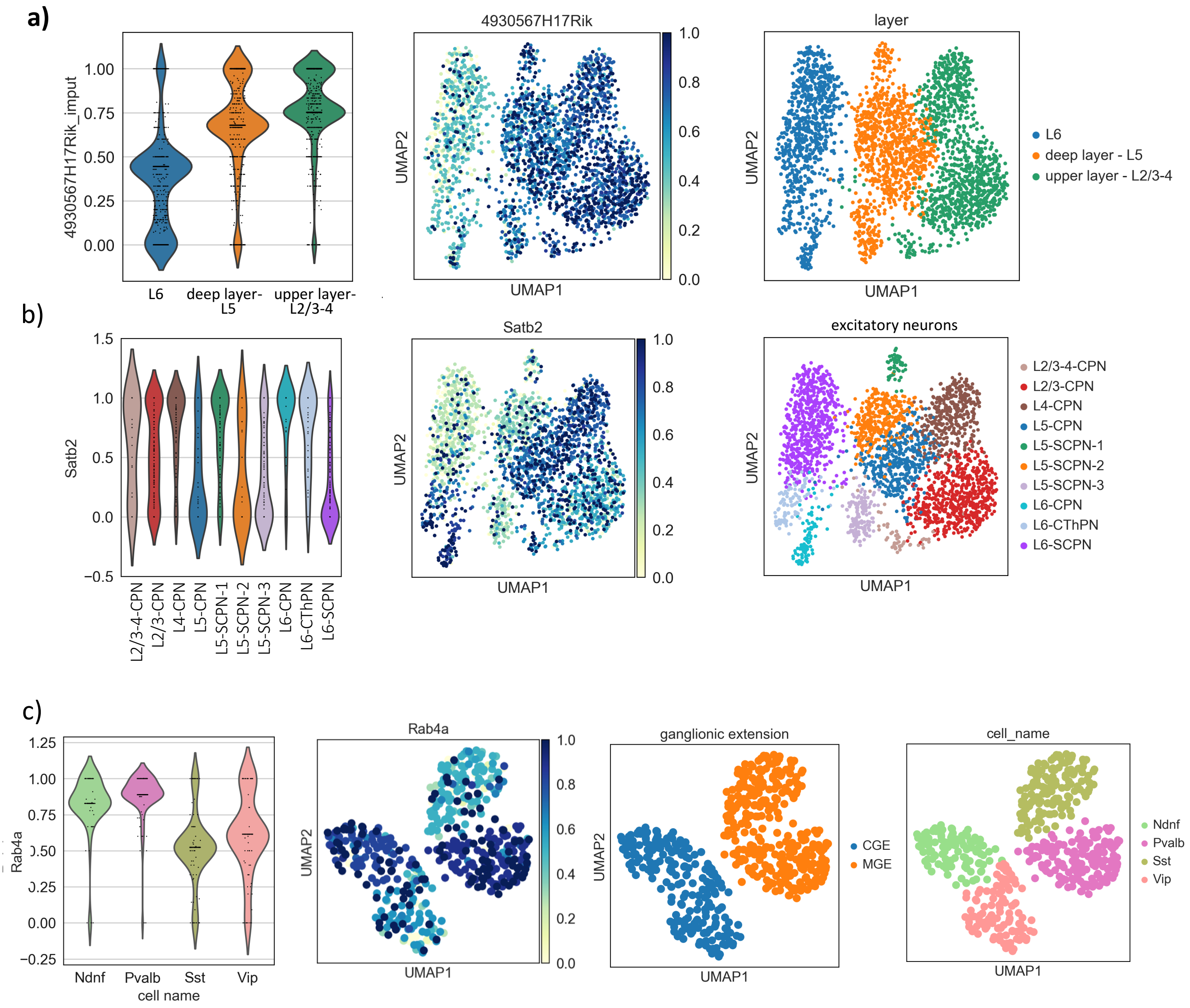


**Suppl. Figure 6: a)** DNA methylation levels at *4930567H1Rik*. **b)** DNA methylation levels at *Satb2*. **c)** DNA methylation levels at *Rab4a*

**Suppl. Figure 7: CH methylation**

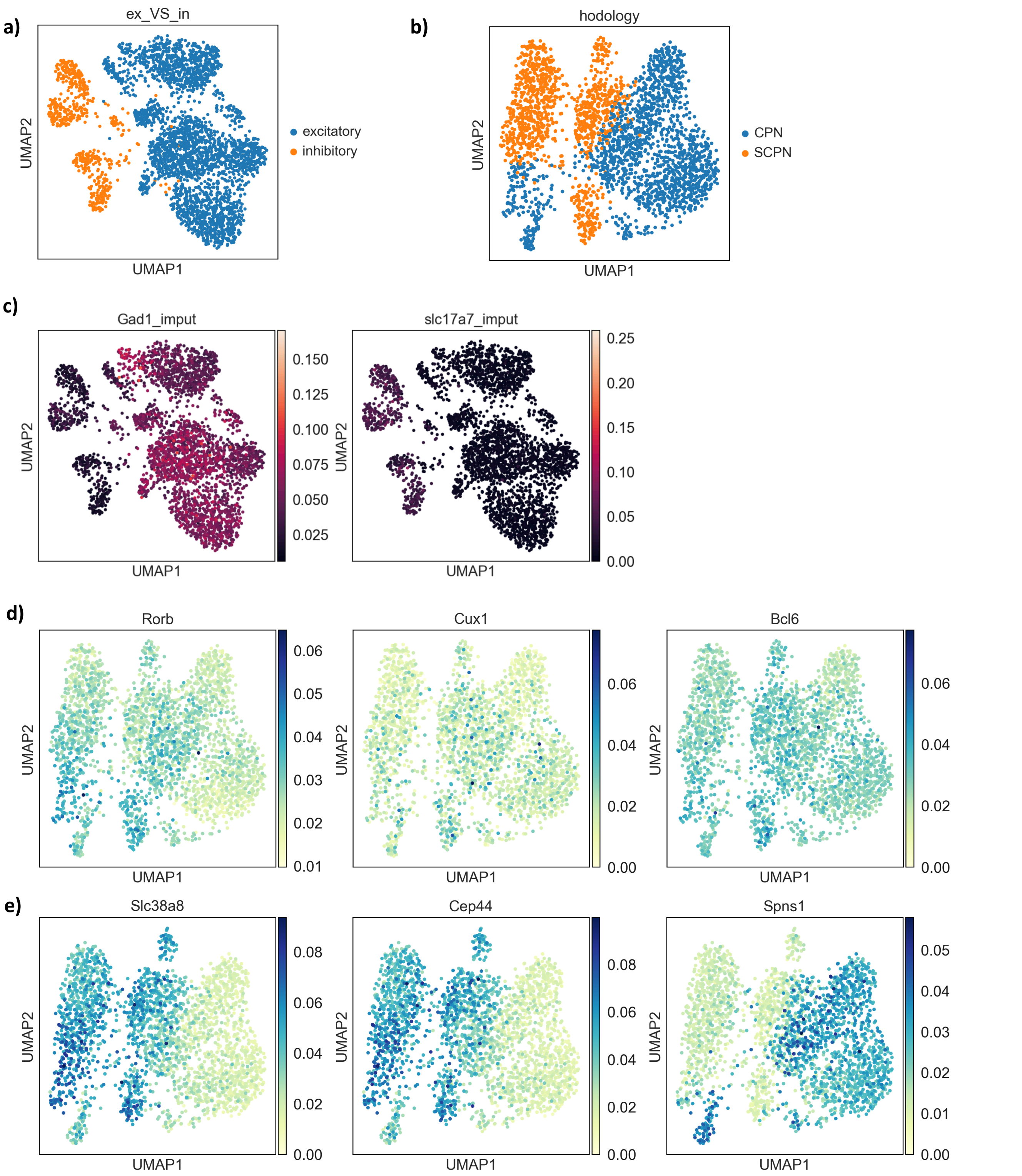


**Suppl. Figure 7: a)** Excitatory and inhibitory neurons (cell type identification based on epiScanpy CG promoter methylation). **b)** CPN vs SCPN excitatory neurons (cell type identification based on epiScanpy CG promoter methylation) **c)** CH methylation levels at canonical markers of inhibitory (Gad1) and excitatory (Slc17a7) neurons. **d)** CH methylation levels at CPN (Rorb and Cux1) and SCPN (Bcl6). **e)** CH methylation levels at some epiScanpy top ranked differentially methylated gene bodies.

### Suppl. Figure 8: scATAC-seq markers

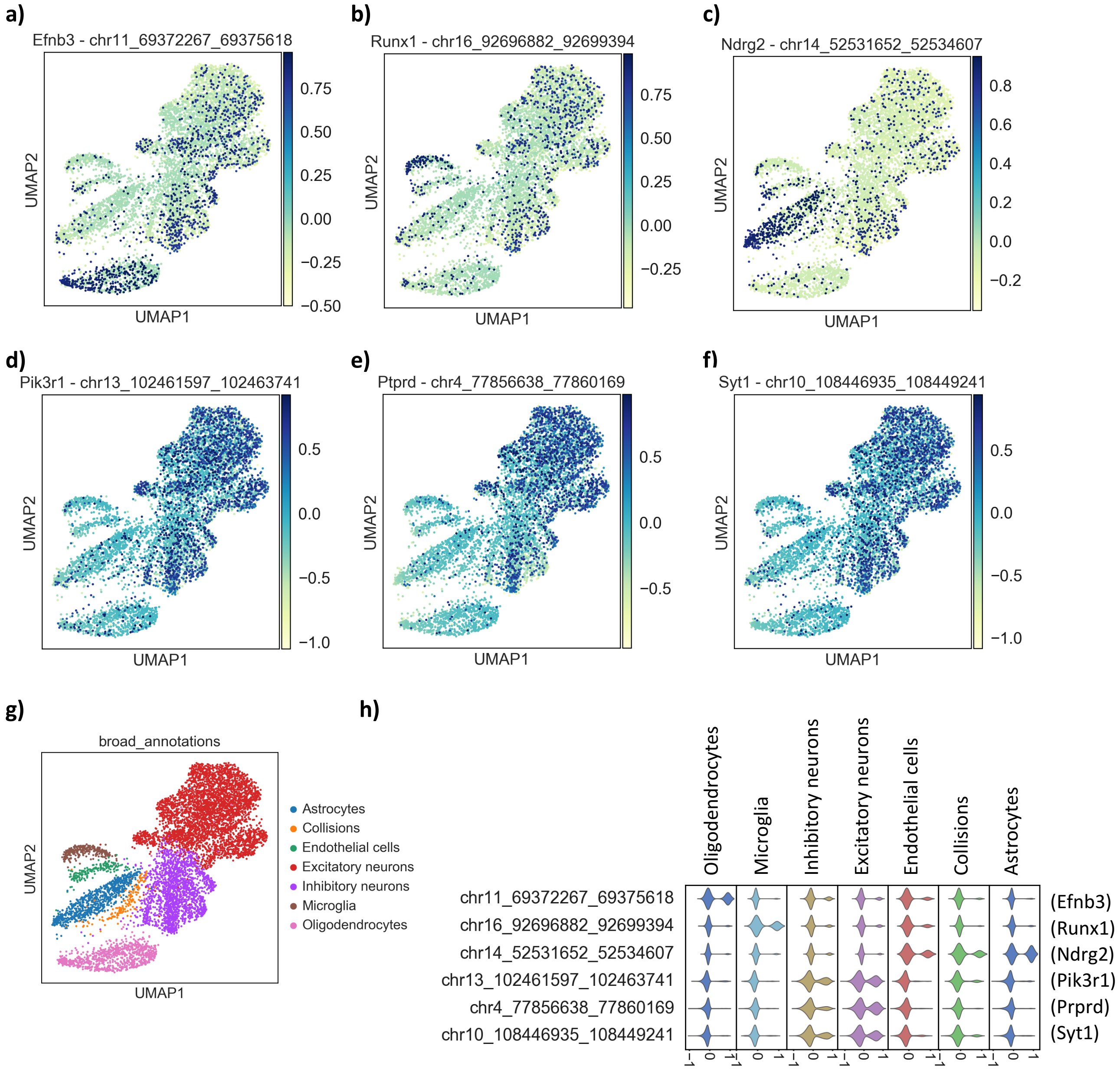


**Suppl. Figure 8:** for the cells in "Prefrontal cortex"+"Whole brain" (8068 cells), openness at the following promoters: **a)** Efnb3 (marker for oligodendrocytes), **b)** Runx1 (marker for microglia), **c)** Ndrg2 (marker for astrocytes), **d)** Pik3r1, **e)** Prprd and **f)** Syt1 (markers for neurons). **g)** UMAP with cell type labels obtained with epiScanpy. **h)** violin plot showing openness at the same markers for the different clusters.

**Suppl. Figure 9: comparison to Luo *et al.* 2017 annotation for scDNAmeth**

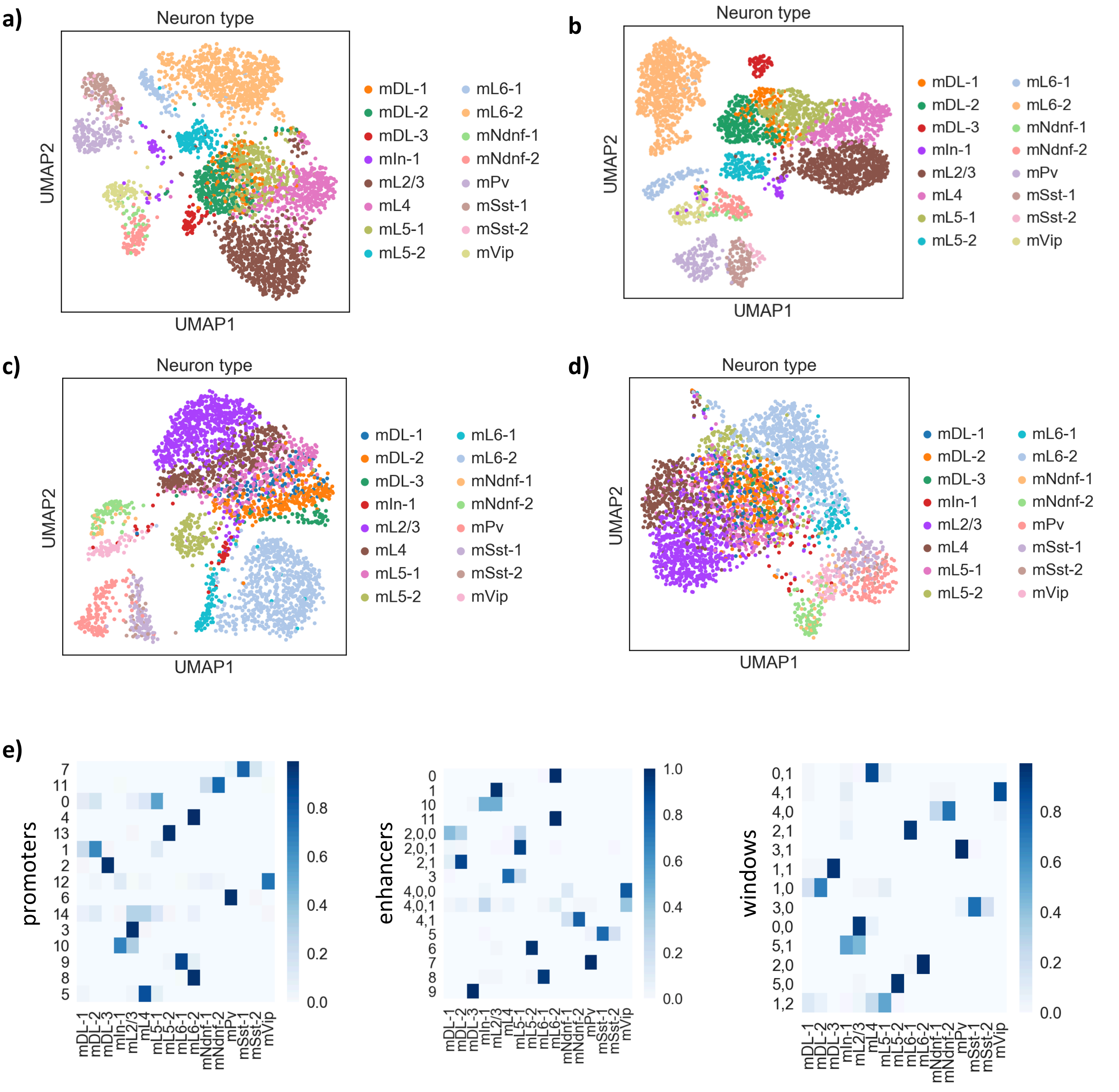


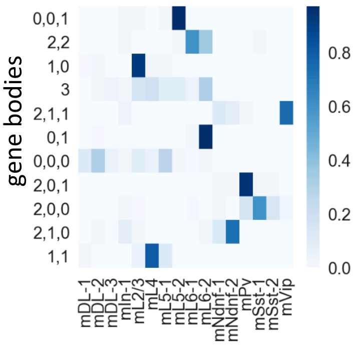


**Suppl. Figure 9:** UMAP visualization with the original annotation from Luo *et al.* (2017) superimposed, for clustering performed using: **a)** promoters, **b)** enhancers, **c)** windows and **d)** gene bodies. **e)** heat map with cell type correspondence between epiScanpy cluster annotation (rows) and the original Luo *et al.* (2017) annotation (columns).

**Suppl. Figure 10: comparison to Cusanovich *et al.* 2018 annotation for scATAC-seq**

a)

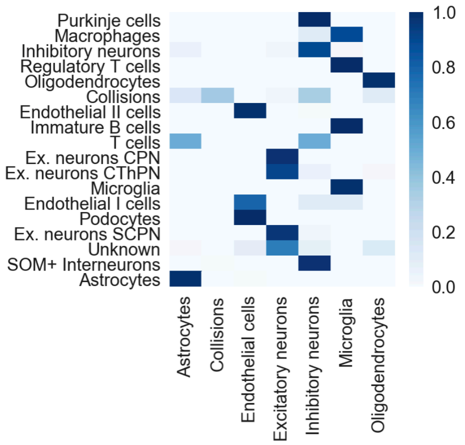

b)

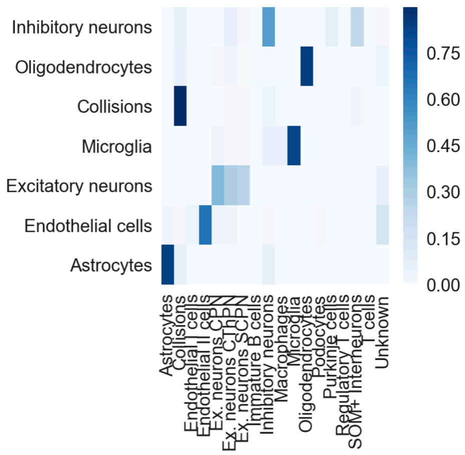

c)

| | Astrocytes | Collisions | Endothelial cells | Excitatory neurons | Inhibitory neurons | Microglia | Oligodendrocytes |
|---|---|---|---|---|---|---|---|
| **Purkinje cells** | 0 | 0 | 0 | 0 | 135 | 0 | 0 |
| **Macrophages** | 0 | 0 | 0 | 0 | 2 | 16 | 0 |
| **Inhibitory neurons** | 60 | 7 | 2 | 23 | 907 | 14 | 1 |
| **Regulatory T cells** | 0 | 0 | 0 | 0 | 0 | 1 | 0 |
| **Oligodendrocytes** | 0 | 0 | 0 | 0 | 3 | 0 | 742 |
| **Collisions** | 58 | 141 | 3 | 11 | 133 | 0 | 49 |
| **Endothelial II cells** | 0 | 0 | 120 | 0 | 1 | 0 | 0 |
| **Immature B cells** | 0 | 0 | 0 | 0 | 0 | 1 | 0 |
| **T cells** | 1 | 0 | 0 | 0 | 1 | 0 | 0 |
| **Ex. neurons CPN** | 0 | 0 | 6 | 1567 | 11 | 7 | 12 |
| **Ex. neurons CThPN** | 1 | 2 | 5 | 1162 | 88 | 3 | 24 |
| **Microglia** | 0 | 0 | 1 | 0 | 1 | 221 | 1 |
| **Endothelial I cells** | 0 | 0 | 8 | 0 | 1 | 1 | 0 |
| **Podocytes** | 0 | 0 | 2 | 0 | 0 | 0 | 0 |
| **Ex. neurons SCPN** | 1 | 2 | 1 | 1055 | 27 | 4 | 8 |
| **Unknown** | 4 | 0 | 27 | 198 | 21 | 1 | 36 |
| **SOM+ Interneurons** | 2 | 4 | 0 | 3 | 434 | 0 | 0 |
| **Astrocytes** | 675 | 1 | 6 | 0 | 2 | 0 | 0 |

### Suppl. Figure 10 (continued)

**d)**

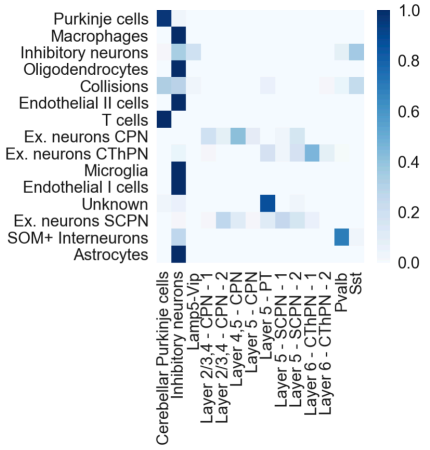


**e)**

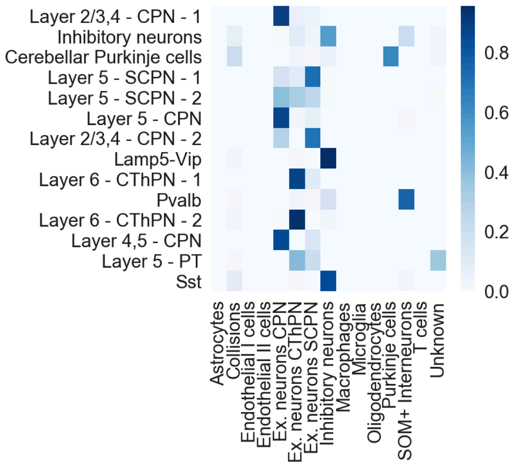


**f)**

| | Cerebellar Purkinje cells | Inhibitory neurons | Lamp5-Vip | Layer 2/3,4 - CPN - 1 | Layer 2/3,4 - CPN - 2 | Layer 4,5 - CPN | Layer 5 - CPN | Layer 5 - PT | Layer 5 - SCPN - 1 | Layer 5 - SCPN - 2 | Layer 6 - CThPN - 1 | Layer 6 - CThPN - 2 | Pvalb | Sst |
|---|---|---|---|---|---|---|---|---|---|---|---|---|---|---|
| **Purkinje cells** | 130 | 4 | 0 | 0 | 0 | 0 | 0 | 0 | 0 | 0 | 0 | 0 | 0 | 1 |
| **Macrophages** | 0 | 2 | 0 | 0 | 0 | 0 | 0 | 0 | 0 | 0 | 0 | 0 | 0 | 0 |
| **Inhibitory neurons** | 13 | 316 | 178 | 1 | 0 | 4 | 0 | 3 | 4 | 6 | 3 | 2 | 68 | 332 |
| **Oligodendrocytes** | 0 | 3 | 0 | 0 | 0 | 0 | 0 | 0 | 0 | 0 | 0 | 0 | 0 | 0 |
| **Collisions** | 46 | 42 | 4 | 0 | 0 | 0 | 0 | 9 | 0 | 0 | 0 | 2 | 7 | 34 |
| **Endothelial II cells** | 0 | 1 | 0 | 0 | 0 | 0 | 0 | 0 | 0 | 0 | 0 | 0 | 0 | 0 |
| **T cells** | 1 | 0 | 0 | 0 | 0 | 0 | 0 | 0 | 0 | 0 | 0 | 0 | 0 | 0 |
| **Ex. neurons CPN** | 1 | 6 | 0 | 318 | 123 | 654 | 142 | 3 | 63 | 262 | 1 | 1 | 2 | 2 |
| **Ex. neurons CThPN** | 6 | 61 | 3 | 21 | 0 | 2 | 6 | 218 | 37 | 213 | 566 | 99 | 12 | 6 |
| **Microglia** | 0 | 1 | 0 | 0 | 0 | 0 | 0 | 0 | 0 | 0 | 0 | 0 | 0 | 0 |
| **Endothelial I cells** | 0 | 1 | 0 | 0 | 0 | 0 | 0 | 0 | 0 | 0 | 0 | 0 | 0 | 0 |
| **Unknown** | 8 | 13 | 0 | 0 | 0 | 0 | 0 | 190 | 1 | 7 | 0 | 0 | 0 | 0 |
| **Ex. neurons SCPN** | 3 | 16 | 0 | 17 | 290 | 108 | 14 | 108 | 274 | 170 | 74 | 0 | 7 | 1 |
| **SOM+ Interneurons** | 0 | 117 | 1 | 0 | 0 | 0 | 2 | 0 | 1 | 0 | 0 | 0 | 301 | 15 |
| **Astrocytes** | 0 | 2 | 0 | 0 | 0 | 0 | 0 | 0 | 0 | 0 | 0 | 0 | 0 | 0 |

**Suppl. Figure 10: a)** Heat map with cell type correspondence between epiScanpy cluster annotation (columns) and the original Cusanovich *et al.* (2018) annotation (rows). **b)** Heat map with cell type correspondence between epiScanpy cluster annotation (rows) and the original Cusanovich *et al.* (2018) annotation (columns). **c)** cell number correspondence between epiScanpy and the original Cusanovich*et al.* (2018) annotation. **d)** Heat map with cell type correspondence for neurons only between epiScanpy cluster annotation (columns) and the original Cusanovich*et al.* (2018) annotation (rows). **e)** Heat map with cell type correspondence for neurons only between epiScanpy cluster annotation (rows) and the original Cusanovich *et al.* (2018) annotation (columns). **f)** cell number correspondence for neurons only between epiScanpy and the original Cusanovich *et al.* (2018) annotation.

### Suppl. Figure 11: Mislabeled cluster markers

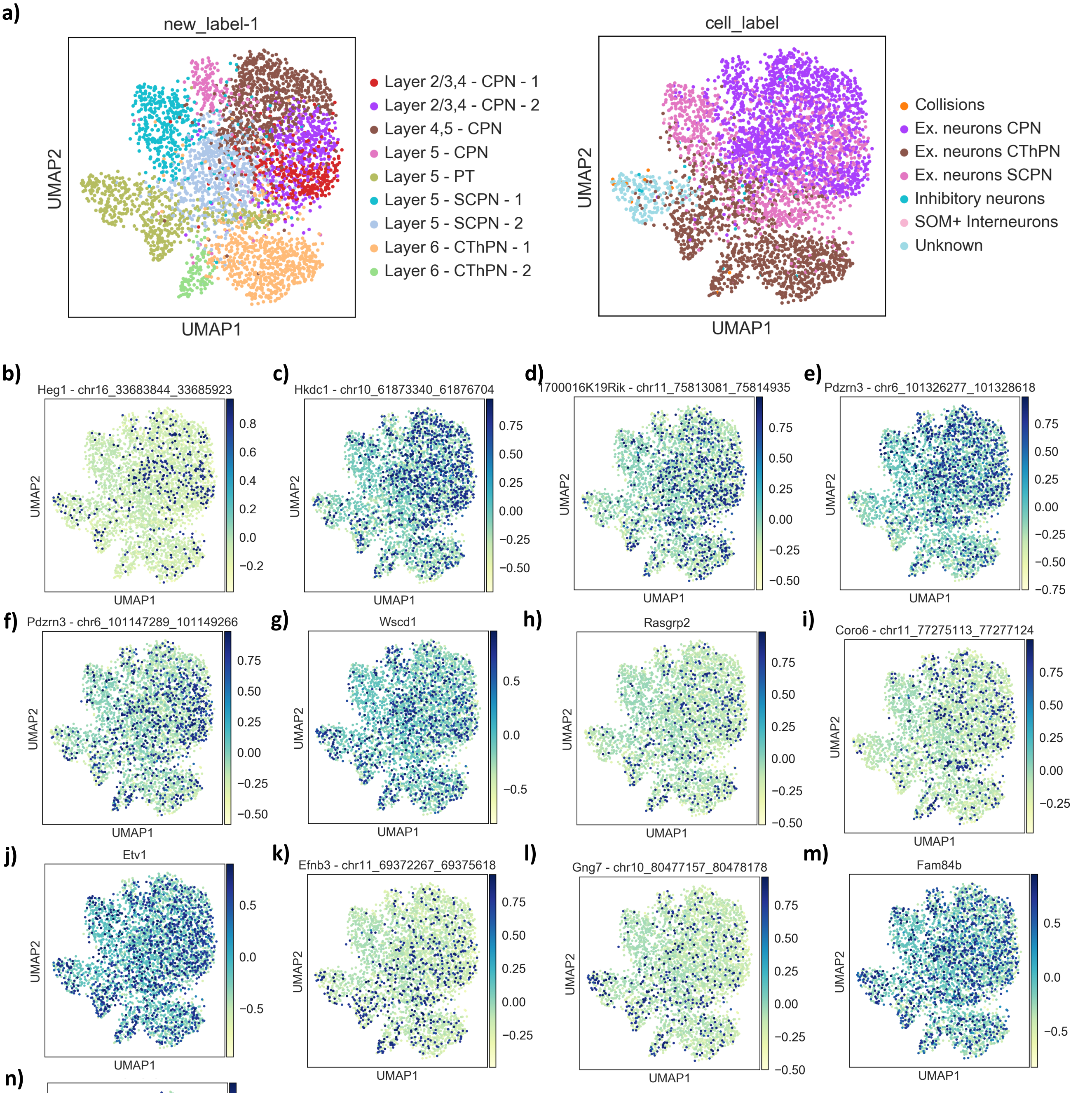


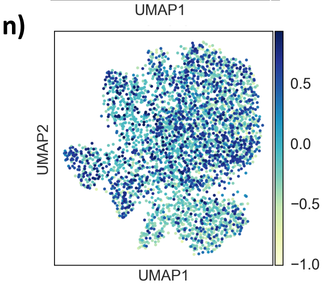


**Suppl. Figure 11: a)** UMAP for neurons with the corresponding epiScanpy label (left) and with the original Cusanovich *et al.* (2018) annotation superimposed. In this low dimensional representation some of the cells originally labelled by Cusanovich *et al.* as “Ex. Neurons CPN” and “Ex. Neurons SCPN” are completely mixed, and in particular the cells labelled “Ex. Neuron SCPN” by Cusanovich *et al.* become labelled “Layer 2/3,4 CPN” by epiScanpy . Openness at several CPN markers for **b-d)** Layer 4 CPN markers and **e-f)** Layer 2/3 CPN markers showing how these markers are open in these cells. Openness at several SCPN markers (**g-k)** for SCPN markers and **l-n**) for subtype specific SCPN markers), showing that SCPN markers are either not open or are open in all cells.

**Suppl. Figure 11 (continued)**

**o)**

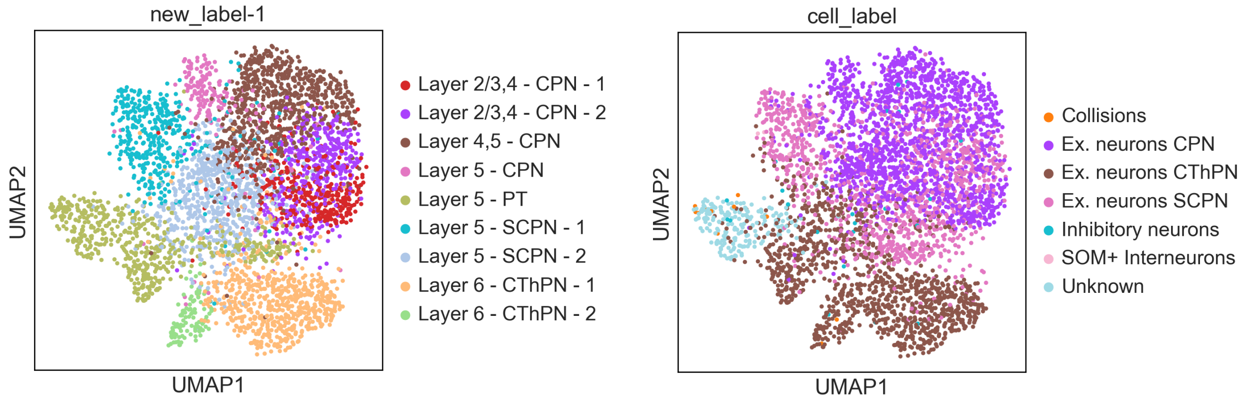


**p)**

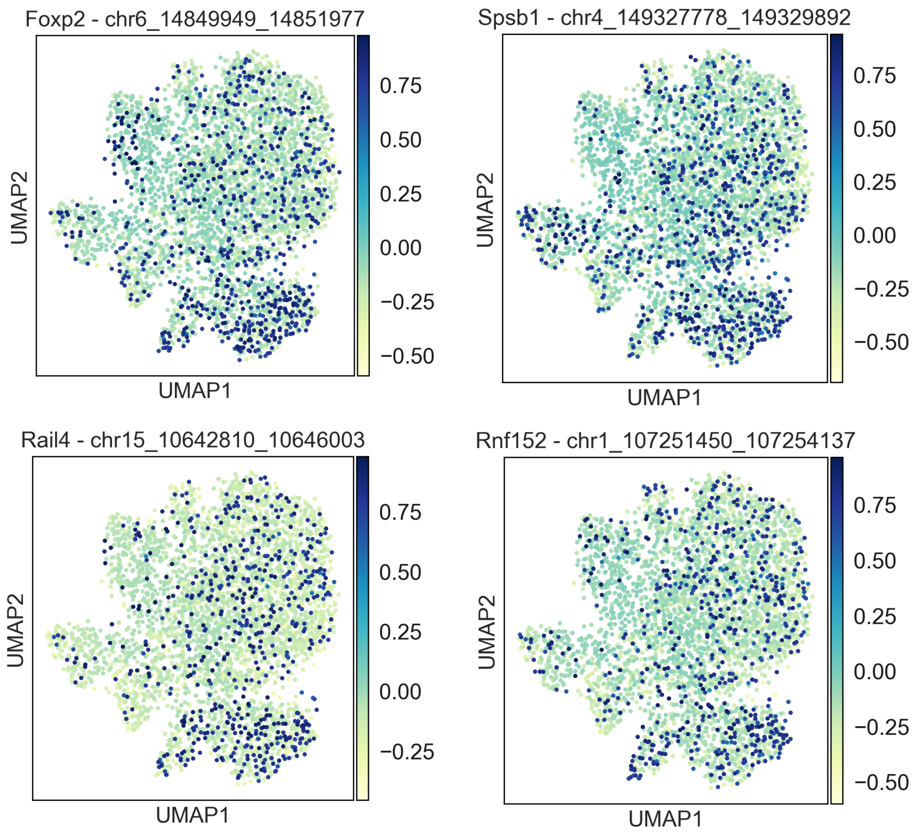


**Suppl. Figure 11 (continued): o)** UMAP for neurons with the corresponding epiScanpy label (left) and with the original Cusanovich *et al.* (2018) annotation superimposed, where one can see that some of the cells originally labelled as “Ex. Neurons CThPN” by Cusanovich *et al.* are labelled as “Layer 5 PT” by epiScanpy. **p)** Openness at Foxp2, Spsb1, Rail4 and Rnf152, all canonical markers for CThPN neurons, showing how they are closed in the cells labelled as “CThPN” by Cusanovich *et al.* 2018 that have been labelled “Layer 5 PT” by epiScanpy.

**Suppl. Figure 12: Markers among multi-omics**

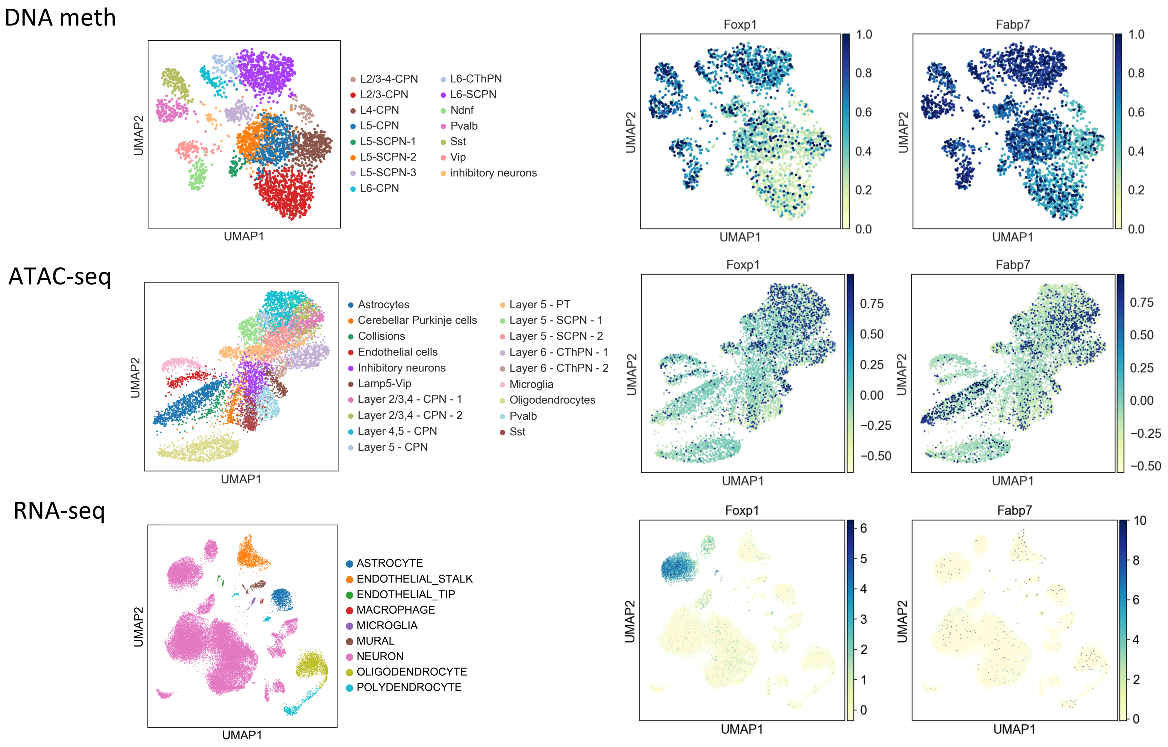


**Suppl. Figure 12:** Example for a shared (Foxp1) and a non-shared (Fabp7) marker between Atlases. In the middle and right panels, the promoter CG methylation (top), promoter openness (middle) and gene expression (bottom) are shown.

**Suppl. Figure 13: scATAC-seq hematopoiesis cell type identification**

**a)**

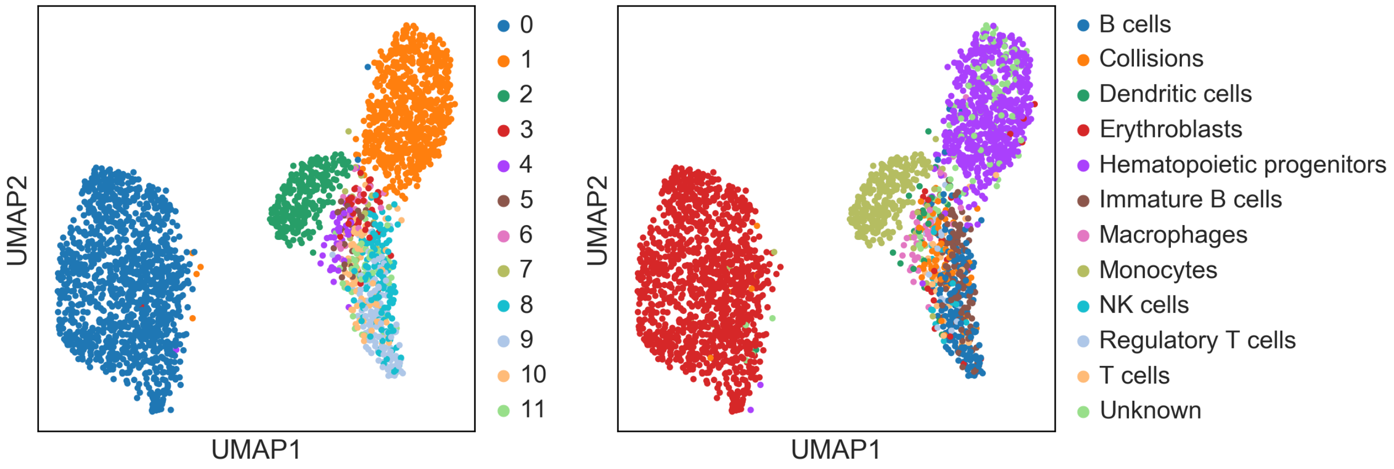


**b)**

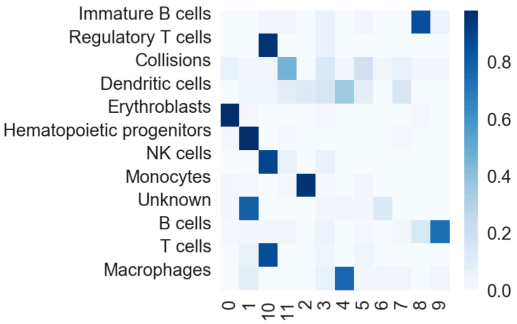


0 – Erythroblasts
1 – Hematopoietic progenitors
2 – Monocytes
3 – Unknown
4 – Macrophages
5 – Collisions
6 – Unknown
7 – Unknown
8 – immatures B cells
9 – B cells
10 – T cells & NK cells
11 – Collisions

**Suppl. Figure 13: a)** Louvain clustering based on peaks for "BoneMarrow" cells (Cusanovich *et al.* 2018) with epiScanpy cluster number (left) and the corresponding cell type annotation from Cusanovich *et al.* 2018 (right). **b)** Cell type correspondence between the two annotations (Cusanovich *et al.* 2018 annotation in rows, epiScanpy Louvain cluster numbers in columns).

**Suppl. Figure 14: 10k PBMC from the chromium 10x scATAC-seq platform**

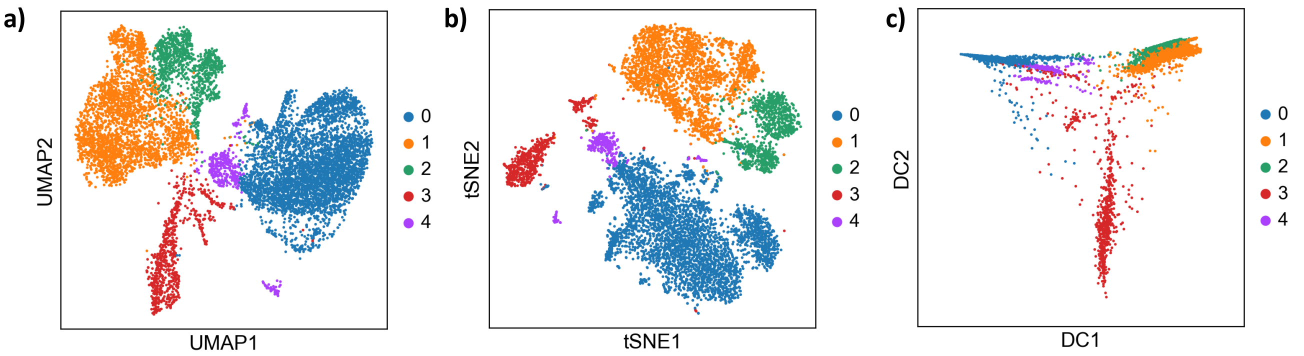


**Suppl. Figure 14: a)** UMAP with Louvain clustering for 10k PBMCs sequenced for scATAC-seq on the 10X platform. **b)** tSNE with Louvain clustering for 10k PBMCs sequenced for scATAC-seq on the 10X platform. **c)** diffusion pseudotime for the same dataset.
